# Primate MHC class I from Genomes

**DOI:** 10.1101/266064

**Authors:** D.N. Olivieri, F. Gambón-Deza

## Abstract

The major histocompatibility complex (MHC) molecule plays a central role in the adaptive immunity of jawed vertebrates. Allelic variations have been studied extensively in some primate species, however a comprehensive description of the number of genes remains incomplete. Here, a bioinformatics program was developed to identify three MHC Class I exons (EX2, EX3 and EX4) from Whole Genome Sequencing (WGS) datasets. With this algorithm, MHC Class I exons sequences were extracted from 30 WGS datasets of primates, representatives of Apes, Old World and New World monkeys and prosimians. There is a high variability in the number of genes between species. From human WGS, six viable genes (HLA-A, -B, -C, -E, -F, and -G) and four pseudogene sequences (HLA-H, -J, -L, -V) are obtained. These genes serve to identify the phylogenetic clades of MHC-I in primates. The results indicate that human clades of HLA-A -B and -C were generated shortly after the separation of Old World monkeys. The clades pertaining to HLA-E, -H and -F are found in all primate families, except in Prosimians. In the clades defined by HLA-G, -L and -J, there are sequences from Old world monkeys. Specific clades are found in the four primate families. The evolution of these genes is consistent with birth and death processes having a high turnover rates.

## 1. Introduction

The Major Histocompatibility Complex (MHC) molecules are amongst the most intensely studied of the adaptive immune system, especially in primates and the mouse since they are responsible for transplant rejection and antigen presentation. These molecules were first discovered in humans on the surface of leukocyte membranes and were given the name HLA (Human Leukocyte Antigen). Although the molecules of MHC were originally studied as the cause of transplant rejection, their primary function is in the defense of the organism against pathogens. The molecule of MHC class I presents endogenous antigen to the CD8+ T lymphocytes and the MHC class II presents exogenous antigens to the CD4+ T lymphocytes. Other molecules related to these pathways are present in the same chromosome MHC regions, sometimes referred to collectively as the MHC class III genes (i.e., complement proteins, cytokines, etc.), although this is a nomenclature that is simply used to make a distinction from the MHCI and MHCII genes.

MHC class I (MHC-I) are encoded by genes with at least 6 exons. The exons of particular interest are exons-2 and -3 (EX2 and EX3, respectively) encoding the protein domains α1 and α2, that are responsible for presenting peptides to TCR. These domains also represent regions of the highest allelic variation. These genes are essential for the innate and adaptive immune responses and are subject to environmental and evolutionary pressures together with likely coevolution effects through their interaction with TCR and KIR (de Groot et al., 2015; Garcia et al., 2009).

In humans, the MHC-I region consists of six genes. The HLA-A, -B and -C genes are highly polymorphic and are expressed by all cells. These molecules are considered the classical MHC-I molecules and their role is to present peptides to cytotoxic T lymphocytes. Unlike the classical MHC genes, the nonclassical genes, HLA-E, -F and -G, exhibit limited polymorphism. In particular, HLA-E is a CD94/NKG2A ligand, HLA-G cells are found only in the trophoblast (Castro et al., 1996; Djurisic & Hviid, 2014; Lynge-Nilsson et al., 2014) and HLA-F is involved in NK cell signaling (Lee et al., 2010). Both HLA-F and HLA-G are expressed by a restricted set of cell populations.

Apart from these MHC-I genes in humans, complete genes have been identified that are not expressed because of different causes: some are pseudogenes (e.g., the known pseudogenes HLA-H, -K, -J and -L) (Moscoso et al., 2006; Heinrichs & Orr, 1990) as well as individual MHC-like exons that are not found in tandem with other valid structural exons (e.g., those possessing stop-codons) needed to form functional MHC-I molecule (Horton et al., 2004).

All humans possess the haplotype with six MHC-I expressed genes. Orthologs to these haplotypes have been sought in Apes, Old World, and New World monkeys. Previous studies have described MHC-I genes orthologous to those in humans, and the human pseudogene, HLA-H, is actually a viable MHCI gene in chimpanzees and gorillas (Wilming et al., 2013). At present, there are detailed descriptions of the MHC loci in Apes, but few such descriptions in evolutionarily more distant primates. For these more distant primate species, several RNA studies have attempted to characterize allelic variability (de Groot et al., 2012). Nonetheless, these sequences have not been studied within the context of the germline MHC genes from all these species.

In recent years, genome sequences of primates have become publicly available in the form of assembled WGS datasets. These assemblies consist of relatively large contigs containing most of the genomic sequences. In this paper, we describe the analysis of MHC-I exon sequences EX2, EX3 and EX4 from primates that were identified from WGS data using a new bioinformatics tool, called `MHCfinder`, freely availble at http://vgenerepertoire.org/. The primate datasets we utilized in this study are indicated in the phylogenetic ordering of 1, obtained from molecular studies Perelman et al. (2011); Rogers & Gibbs (2014) and divergence times further confirmed with sets of published works summarized by TimeTree (Hedges et al., 2006). From the sequences found by `MHCfinder`, this wealth of detailed genomic information may help to clarify evolutionary processes that have shaped the MHC-I genes in primates.

**Figure 1:**
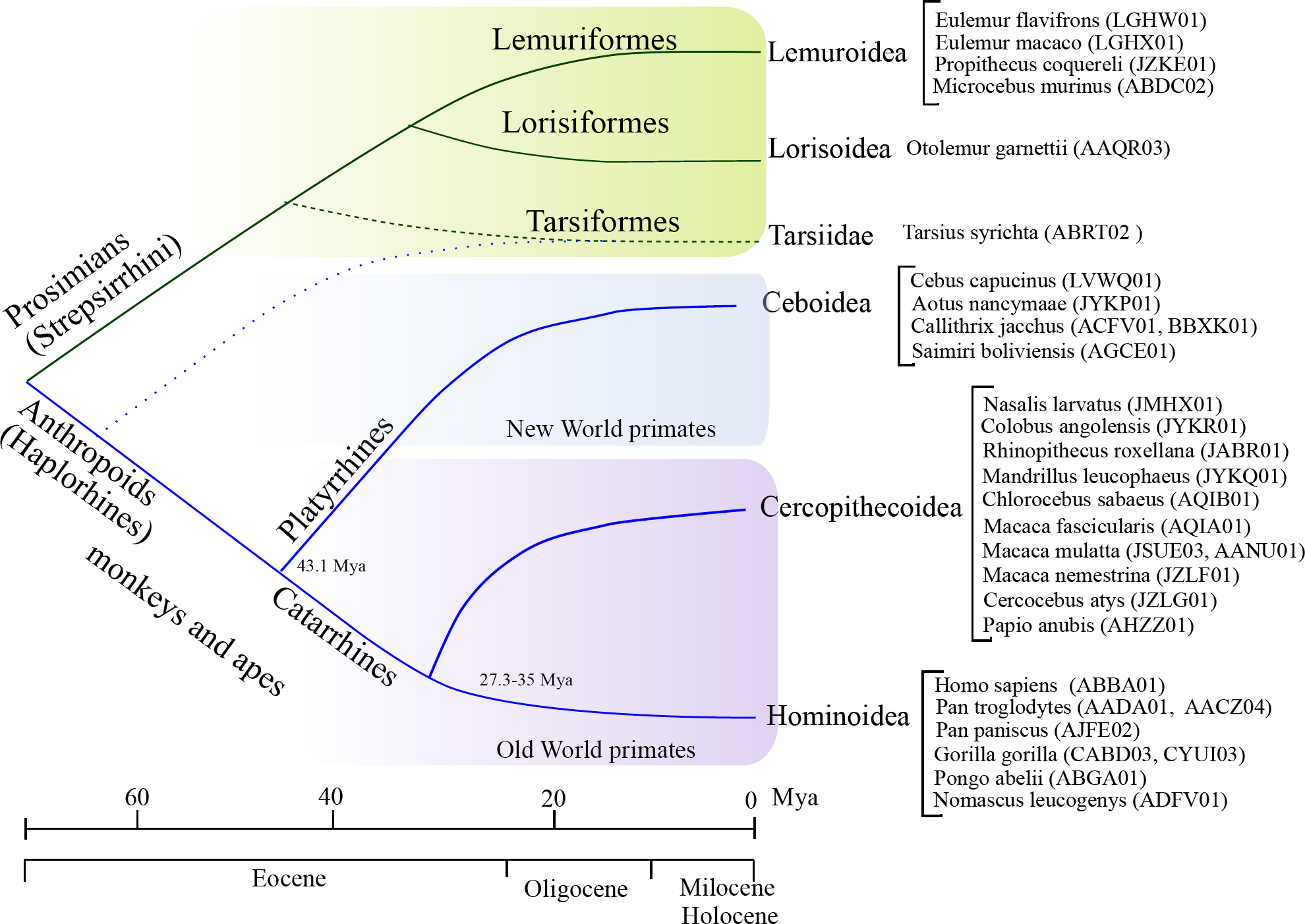
The phylogenetic tree of the primates indicating the species studied in this work. The tree is based upon divergence times obtained from (Hedges et al., 2006), and previous molecular phylogenetic studies Perelman et al. (2011); Rogers & Gibbs (2014).

## 2. Methods

### 2.1. Datasets

The WGS assembly datasets of 30 primate species were obtained from the NCBI in the form of FASTA files consisting of assembled contigs, or for more mature projects, scaffolds and/or fully constructed chromosomes. The average genome coverage in these datasets is > 15−20× with contig assembly N50 > 15kbs. A detailed summary of the accession numbers and relevant assembly parameters can be found in Table 1.

### 2.2. Software

While there are six viable MHC-I genes in *Homo sapiens*, the number of MHC-I genes varies considerably across other mammal species. Nonetheless, the MHC-I exon architecture is nearly universal across all mammal species (Birch et al., 2006). A diagram of the *Homo sapiens* MHC-I (HLA) exon structure and corresponding protein domains was described (Lefranc et al., 2005) (see also sequence repositories of the by the IMGT) and the a large collection of MHC-I alleles at the IPD-MHC database (Robinson et al., 2011). The relevant genomic signals (i.e., the exons, introns, 5′-URT and 3′-URT) can be identified to a high degree of accuracy without resorting to general gene finding software tools. Therefore, viable exons (i.e., those exons that could form a functionally expressed MHC-I molecule) can be accurately determined using homology criteria with a supervised machine learning classifier.

The `MHCfinder` program extracts exons EX2 and EX3 that encode the α1 and α2 domains, respectively, together with exon EX4 that encodes the constant domain. Although other exons constitute the full MHCI gene, these three exons are of particular interest for characterizing and comparing MHCI genes within and amongst species. Thus, `MHCfinder` ignores the peptide leader (L) (given by EX1), as well as all peptides corresponding to the transmembrane and cytoplasmic (*Tm*, indicated by EX5 through EX8). While the exon/intron structure of MHC Class I is thought to be universal across jawed vertebrates, the specific intron spacing between the exon sequences EX2, EX3, and EX4 varies considerably. As such, the algorithm only imposes a simple structural requirement that EX2, EX3 and EX4 are found in a tandem arrangement along the DNA sequence, but places no hard restrictions on the intron separation.

Figure 2 summarizes the principal steps of the `MHCfinder` algorithm. This program was implemented as a multi-threaded application in the python programming language, with the biopython library (Cock et al., 2009) for low-level sequence analysis, and the scikits library (Pedregosa et al., 2011) for machine learning tasks. First, a Tblastn query from a consensus protein sequences from known MHC-I exons (EX2, EX3, and EX4 from humans) is made against all available primate WGS datasets. The search result is a listing of candidate WGS contigs likely to contain valid exons, together with the position of the matching nucleotide sequence and similarity scores; this listing is referred to as a *hit table*. The algorithm processes each line of the *hit table*, analyzing a nucleotide region larger than the nucleotide positions in the hit, so as to determine the precise start/stop positions of the exon reading-frame (defined by AG and GT motifs, respectively). Once the exons are identified, they are translated into amino acid sequences by checking all valid reading frames. Those sequences containing stop-codons in the reading frame are discarded, while valid exons are saved and converted into numerical feature vectors (i.e., a unique array of numbers, that uniquely characterizes the string of amino acids). A simple transformation of AA to feature vector was used (based upon the frequency of each AA and pairs of AA), because it was found to discriminate sequences better than other more sophisticated transformation procedures (e.g., those based upon positional physicochemical properties of each AA within the sequence).

From the feature vector representation, a machine learning procedure with a Random Forest (Breiman, 2001) was used to classify the sequence into one of the exon types: EX2, EX3, and EX4. Supervised training is performed by defining these classes with sets of annotated exons from *H. sapiens* and defining a null set from a random background sequences. Binary classification is carried out for each exon type with a background/signal ratio of 3:1, determined empirically. The prediction precision is improved by multiple training/prediction iterations; positively identified sequences are included in the training set for subsequent training/predictions. This process is referred to as iterative supervised learning, and is a common machine learning technique whereby new information is continually accrued to the knowledge base for improving prediction accuracy (e.g., modern speech recognition has benefited from such techniques).

In our gene finding algorithm, `MHCfinder`, a probable functionally expressed MHC-I gene that must contain a tandem arrangement of the three viable exons (i.e. those that do not contain stop codons in the reading frame), EX2-EX3-EX4 along the germline sequence. Nonetheless, viable exons, which are homologous to MHC-I constituent exons, exist in the genome that do not form tandem arrangements (i.e., they are isolated or an exon is missing), and thus, do not express MHC-I molecules. In humans, we found 31 exons, 18 of which are constituents of six functional MHC-I genes; the other exons while viable, must be pseudo-genes. Similar results are seen in all the other species studied.

**Figure 2:**
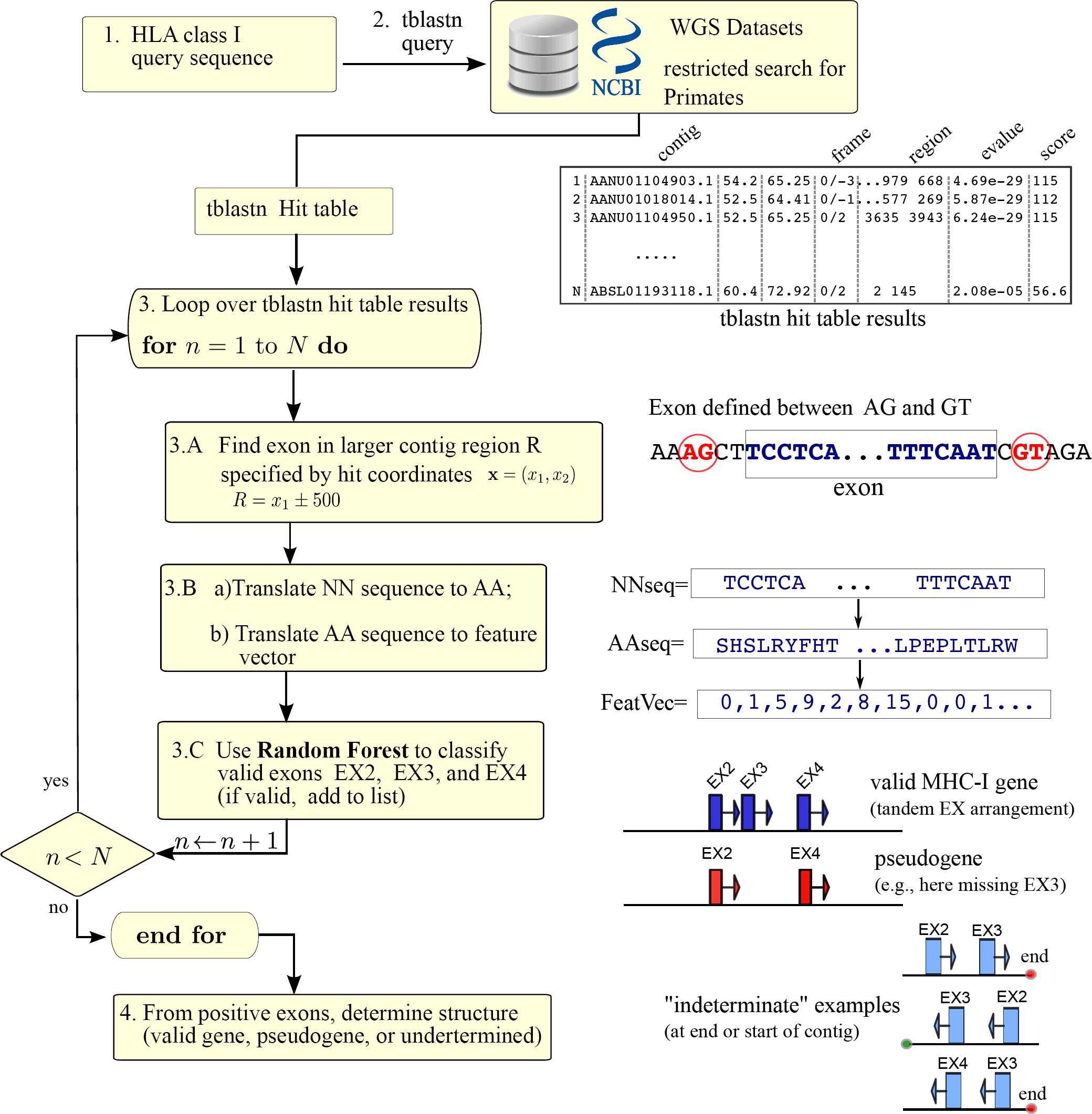
The steps in the MHC-I prediction algorithm. The selection of valid MHC-I exons is based upon a Tblastn pre-selection, an exon reading-frame identification procedure, and classification with a random forest method.

*Tree construction.* To study the phylogenetic relationships from the MHC-I exons, we constructed a large phylogenetic tree by aligning sequences with ClustalO and then used phyML with the WAG matrix (part of the Fast-tree software (Price et al., 2010)). In all cases, 500 bootstrapped samples were made. While trees were studied using the three exon sequences (EX2-EX3-EX4), the final analysis was made only with EX2, since this exon provides the most discriminatory information.

## 3. Results

Exon sequences EX2, EX3 and EX4 of MHC-I were obtained from 30 WGS primate datasets using our software tool, `MHCfinder`, described in the Methods section. These sequences are homologous to the human MHCI and were found by an iterative supervised learning procedure (Methods). As described, these sequences are flanked by splicing signals AG/GT and have an ORF starting two nucleotides after the AG and terminating one nucleotide before the last GT. Exons are considered correct if these conditions are met, while exons possessing stop codons within the reading frame are discarded. Those exons found in a tandem arrangement (i.e. with EX2-EX3EX4 and with nominal intron spacing) are considered candidate MHC-I genes (referred to as *probably viable genes* throughout the rest of the paper, since they have the necessary conditions to be expressed). Valid exons that do not participate in a tandem arrangement are considered probable pseudogenes (referred as *pseudogenes* throughout) or indeterminate if they are found at the extreme ends of the contigs (referred to throughout the text as *indeterminate*). Figure 3 shows graphical maps of the MHC-I exons found in contigs of *G. gorilla*. The exons found in tandem configurations, EX2-EX3-EX4, are indicated as probably viable MHC-I genes. Table 1 lists the number of exons and candidate viable genes per species found from different WGS datasets. Differences in the number of exons between WGS datasets from the same species is indicative of the variability between individuals of the same species, as well as maturity/completeness and sequencing methods used in constructing the assemblies.

At present, there are 42 WGS of *Homo sapiens* in the NCBI repository. From these datasets, `MHCfinder` was used to find all MHC-I exon sequences and the phylogenetic tree of Figure 4 was constructed using the deduced amino acid sequences of EX2. The program tags the EX2 sequence as belonging to one of the following categories: (1) a probable expressed MHC-1 gene (since it is part of a tandem arrangement EX2-EX3-EX4), (2) a pseudogene (because it lacks either EX3, EX4 or both), or (3) an indeterminate gene (because it is found at the extreme edge the contig, but could form a viable gene; see graphic of Figure 2). In the resulting tree, six viable genes (HLA-A, -B, -C, -E, -F and -G) and four pseudogenes (HLA-H, -J, -L and -V) can be discerned. Also, the tree demonstrates that considerable variability exists amongst the classic genes (A, B, and C), forming several lineages. However, with respect to the nonclassical genes (E, F, and G), the sequences are invariable. Also, the known pseudogene sequences, L, J and V are conserved, while the H pseudogene form separate lineages; this may be related to its proximity to the HLA-A locus (Grimsley et al., 1998).

Hominidae diversified approximately 20 (Million years ago) Mya. Eight WGS datasets were used to study the MHC-I exons from the Ape family; the number of exons identified from the WGS of Ape species is provided in Table 1. The number of probable viable genes per species is between two and eight. The presence of pseudogenes is demonstrated by the existence of more exons than needed for the viable genes.

**Table 1:**
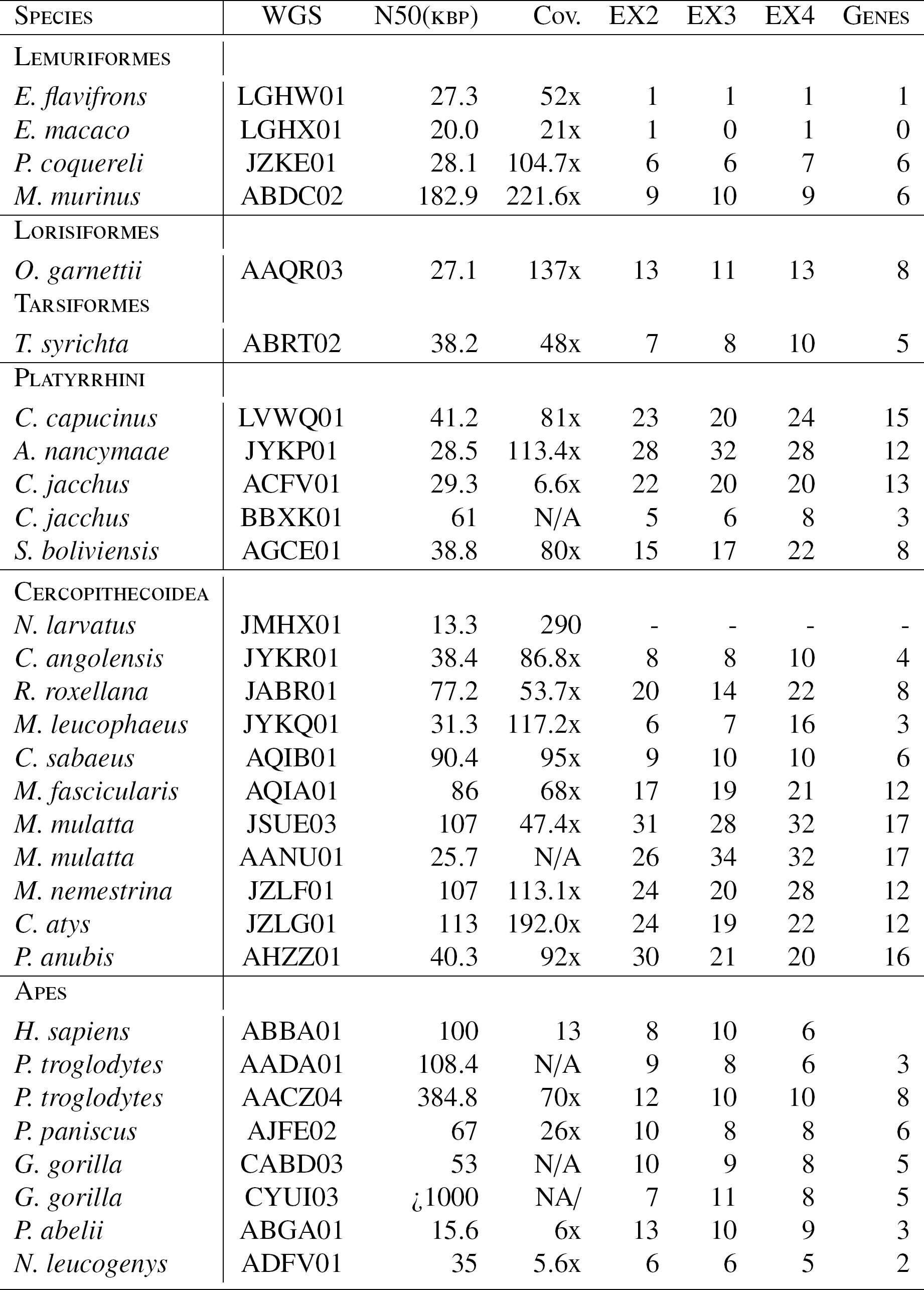
MHC classI Exons and genes of Primates. All values for the contig N50 were > 15kbp, except *N. larvatus* (for which no exons were found). For some assemblies, a reliable number for the coverage could not be ascertained, and are indicated by N/A (not available).

**Figure 3:**
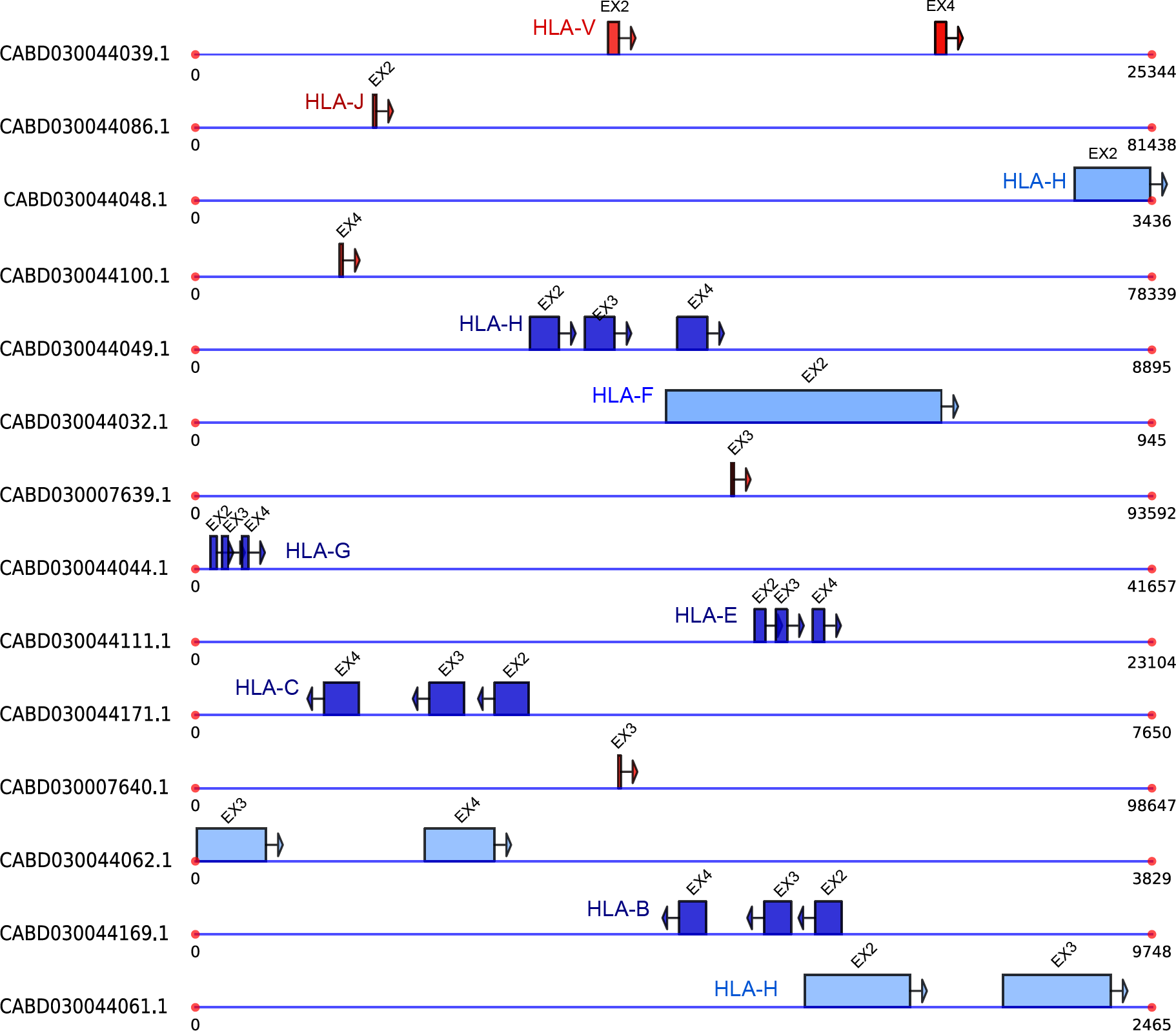
Graphical representation of the MHC-I exons obtained in *G. gorilla* (CABD03) with the `MHCfinder` program. Each contig is represented by a line and exons by boxes. Tandem exon arrangements (EX2-EX3-EX4) are colored blue, indicating the high possibility of being a viable and functionally expressed MHC-I gene (explained in text). Exons not part of tandem arrangements, thus indicative of pseudogenes, are marked in red. Exons that are found at the extreme ends of the contigs may be considered *indeterminate* if they could form tandem arrangements with exons in other contigs (colored in light blue).

The Cercopithecus family consists of monkeys closest to hominids and are also referred to as Old World monkeys. The divergence time between Cercopithecidae and Hominidae is approximately 29Mya. The number of MHC-I genes in these animals varies considerably; between four in *C. angolensis* and 18 in *M. Mulatta* (22 were found in a detailed study (Daza-Vamenta et al., 2004)) (Table 1). In general, the Cercopithecus species possess more MHC-I genes than found in Hominidae species.

The Platyrrhines are monkeys of South and Central America. These species diverged from hominids approximately 43 Mya. Five WGS datasets of Platyrrhines were analyzed to identify MHC-I genes and pseudogenes using `MHCfinder`. The number of viable MHC-I genes found is between nine (in *S. boliviensis*) and 15 (in *C. capucinus*). Lemuriformes and Tarsiiformes represent the oldest extant primates that diverged from other primates approximately 76 Mya. Compared to other taxa of this study, few exons were found from the datasets of *Eulemur* species, perhaps because these sequenced assemblies are still in a nascent stage.

**Figure 4:**
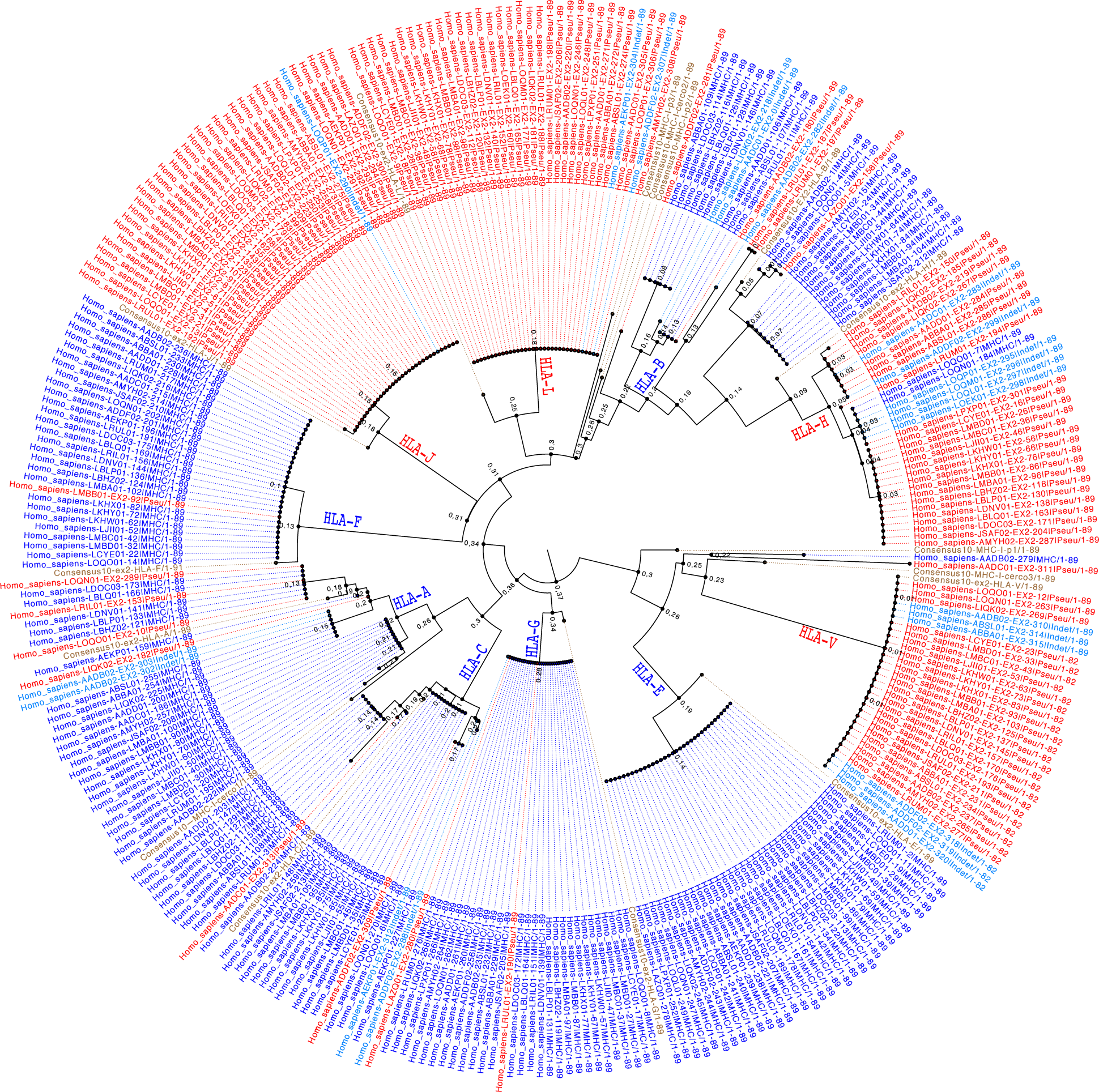
The phylogenetic tree of 333 EX2 sequences of MHC-I found with our |MHCfinder— algorithm from the 42 WGS datasets of *Homo sapiens*. The probable viable genes are colored in blue, pseudogenes are colored red, and those genes that are indeterminate are colored light blue. The sequences in brown correspond to consensus sequences that are used to identify clades. The nodes of the principal clades have been labeled. The tree was constructed after aligning sequences with ClustalO and using the phyML (part of the Fasttree software (Price et al., 2010)) with the gamma parameter, WGS matrix, and 500 bootstrapped samples.

Using `MHCfinder`, 20 WGS datasets from the Simians family (not including humans) were studied, from which 210 viable genes were found. Of the 3 MHC-I exons studied (i.e., EX2, EX3 and EX4), EX2 contain the most information to discriminate genes for classification. A phylogenetic tree (Figure 5) was constructed from 410 EX2 exons found from the primate WGS datasets, consisting of both viable (those having exons in tandem) and nonviable genes. In the tree, clades were collapsed to improve the visualization of results. Human sequences from the HLA isotypes HLA-A, -B, -C, -E, -F, -G, -H, -J, -V and -L genes were aligned with the sequences from non-human primates in order to associate and identify clades homologous to those in humans. Apart from these homologous human clades, other clades were also identified. Three clades containing only sequences from Cercopithecus species are designated with *MHCI-cerco-1, MHC-I-cerco-2* and *MHC-I-cerco 3*. Another three clades with sequences exclusive to Platyrrhini also are identified, indicated by *MHC-I-platy-1, MHC-I-platy-2* and *MHC-I-platy-3*. Also, there are three other clades that have sequences from at least two primate orders and have been labeled *MHC-Ip1, MHC-I-p2* and *MHC-I-p3*. The MHC-Ip1 clade contains of the largest number of sequences, with representatives from all primate species except Hominids, perhaps suggesting a concrete function that was made superfluous in the differentiation of Hominid species.

**Figure 5:**
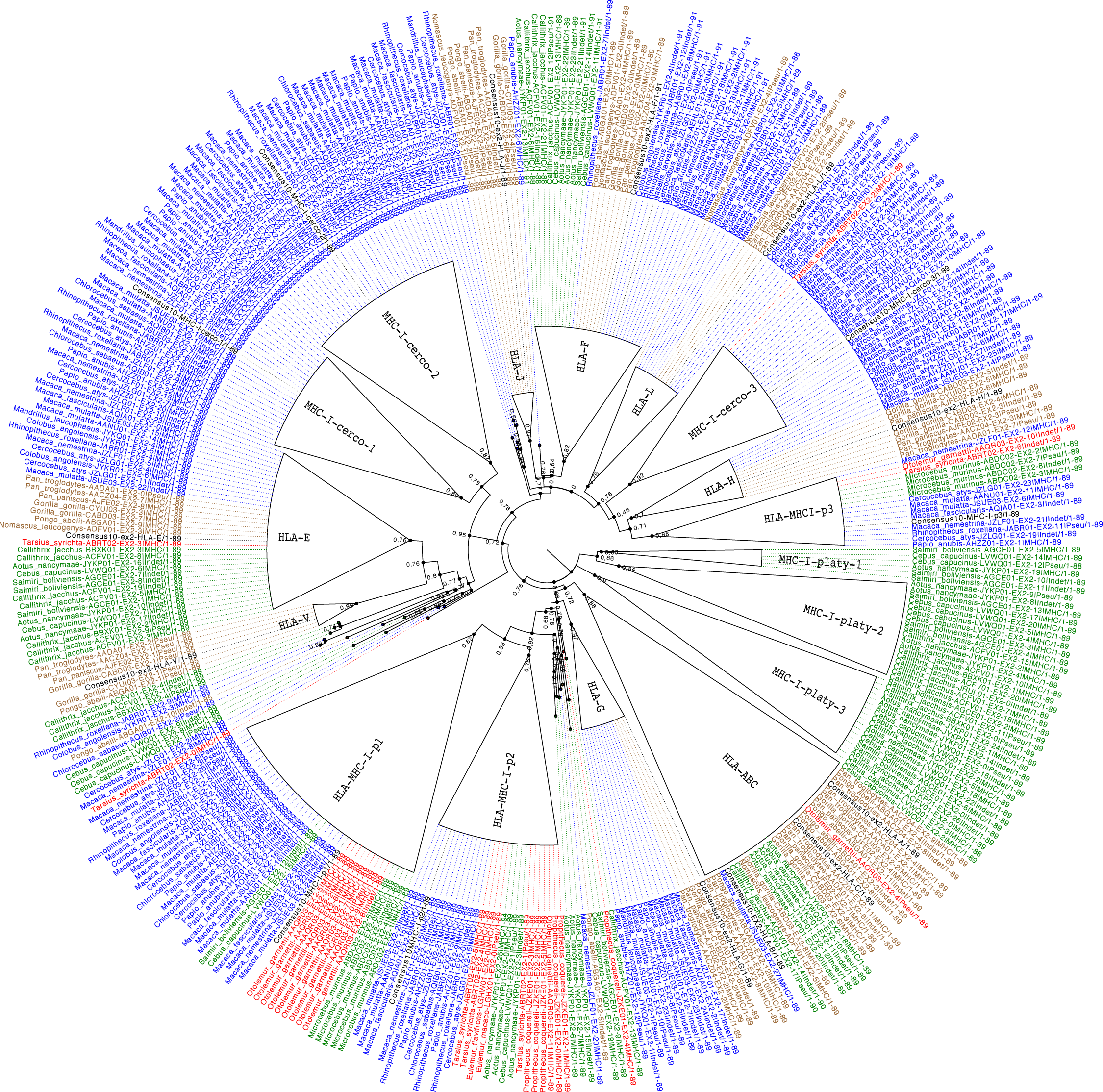
The phylogenetic tree of 417 EX2 sequences of MHC-I found in primate (Prosimians, Platyrrhini, Cercopithecus and Hominidae) WGS datasets with the software `MHCfinder`. Primate orders are distinguished by color codes, indicated in the legend. Clades that are suggestive of representing a gene or group of genes have been collapsed. Each collapsed clade are labeled with the human isoform orthologs, however, those clades not containing human genes are assigned new names (e.g., MHC I-p1, etc.). The nodes of the principal clades have been labeled. The tree was constructed after aligning sequences with ClustalO and using the phyML (part of the Fasttree software (Price et al., 2010)) with the gamma parameter, WGS matrix, and 500 bootstrapped samples. The consensus sequences for identifying clades are marked in black.

Most of the sequences from the HLA-ABC clade (i.e. consisting of HLA-A, -B and -C sequences) are from Hominid species. Also, this clade contains a single sequence from Cercopithecus and seven sequences from Platyrrhinus. HLA A, -B and -C have a common origin and must have been generated after the separation of the Hominids from the Cercopithecus, previously described by other authors (Piontkivska & Nei, 2003). The sequences from these clades constitute the classical HLA-Class I genes. Therefore, it is likely that Cercopithecus and Prosimian species developed other genes that overtook the basic functions of these classical HLA-class I found in Hominids.

From the tree of Figure 5, the HLA-E and -F clades contain sequences from all the primate orders except Prosimians. HLA-G, -H, -J and -L clades have sequences from Apes and Cercopithecus. With the analysis of `MHCfinder`, most of EX2 sequences of Cercopithecus of the HLA-G clade are identified as pseudo-genes, consistent with experimental results (Castro et al., 1996). All the sequences of the HLA-V clade are pseudogenes and are only found in Hominids.

Three clades (indicated by the name *cerco*) are found that contain sequences only from Cercopithecus species. These sequences were generated after the separation of Old World monkeys from the Hominidae family. In a similar way, three clades contain sequences only from Platyrrhine species (indicated by name *platy*.

Another three clades exist that are composed of sequences from at least two primate families, but lacking sequences from Hominids. Of these, the *MHC-I-p1* clade (where *p* is used to indicate primate) is the largest. In particular, the *MHC-I-p1* and *MHCI-p2* clades contain sequences from Prosimians, Platyrrhines and Cercopithecus, while the third clade of this group lacks sequences from Cercopithecus.

### 3.1. Allele assignment to clades

Several studies concerning the gene alleles of MHC-I have been described in non-human primates and are available at the IPD-MHC database (www.ebi.ac.uk/ipd/mhc) (Robinson et al., 2013; de Groot et al., 2012). These alleles were aligned with the germline sequences to classify alleles as classical MHC-I (with allelic variation) or nonclassical MHC-I (without allelic variation).

*M. Mulatta* has been extensively studied to reveal such allelic variation of MHC-I. We obtained allele sequences of *M. Mulatta* from the IPD-MHC database and aligned them to the germline sequences found by `MHCfinder`. Figure 6 shows the resulting phylogenetic tree. To reduce sequence redundancy, only one representative allele from each allele lineage was included in the tree construction. The Mamu-A alleles belongs to the genes that define the MHC-I-p2 and MHC-I-cerco-3 clades. In previous publications, difficulties have been described in assigning specific orthology to the Mamu-B alleles (Liu et al., 2013). The tree of Figure 6 resolves this difficulty by showing that Mamu-B alleles correspond to genes belonging to the MHC-I-p1, MHC-I-cerco-1, -cerco2 and HLA-H clades.

*C. jacchus* is considered the reference organism for studies of allelic variation of MHCI genes in New World monkeys. Many published studies (Cao et al., 2015; van der Wiel et al., 2013; Kono et al., 2014) have concluded that the MHC-I alleles in *C. Jacchus* are orthologous to the HLA-G gene. Figure 6 shows the phylogenetic tree of germline EX2 sequences obtained from `MHCfinder` together with allele sequences from the IPDMHC database. The tree confirms the results of these studies, since the majority of alleles are from gene *C. jacchus-ACFV01-13—MHC* belonging to the HLA-G clade. However, the alleles of the lineage *Caja-G*18* belong to the *MHC-I-platy-3* clade.

## 4. Discussion

A bioinformatics program, `MHCfinder` (freely available at http://vgenerepertoire.org/), was developed that identifies the MHC-I exons (EX2, EX3 and EX4) from WGS datasets. The algorithm determines the in-frame exon nucleotide sequences, deduces the amino acid sequence, and transforms it to a 400 element feature vector. Each element of the feature vector represents the frequency of occurrence of each AA and pairs of AA (i.e., one of the 20 × 20 possibilities) within the protein sequence. A supervised learning procedure is used to train a machine learning classifier to recognize exons homologous to those known in humans. We found that feature vectors based upon the simple frequency of AA occurrence transform has a higher classification accuracy (attaining predictive precision of 98%) than other more sophisticated transforms based upon positional physicochemical properties. If the exon architecture is known, small modifications could render the algorithm a more general gene finding method, thereby facilitating rapid identification of gene/specific exons from any species whose genome has been sequenced.

With `MHCfinder`, exon sequences of MHC-I were obtained from 30 primate WGS datasets. The sequences have been made available in the repository vgenerepertoire.org. The program identifies individual exons, referred to here as *viable*, meaning that they have a valid reading frame. If the exons are found with the canonical MHC-I exon/intron structure (i.e., they are arranged in tandem, EX2-EX3-EX4 within the same WGS contig and have a valid intron spacing), then `MHCfinder` considers this a viable gene (i.e., that is likely functionally expressed). Nonetheless, problems associated with WGS datasets (e.g., incorrect sequence assembly or low coverage of certain regions) may result in an underestimate of the total number of viable MHC-I genes found by `MHCfinder`. In WGS datasets with relatively high coverage and N50¿15k (which is the case for most datasets we used), the results of `MHCfinder` agreed with annotated gene results, when available. While the use of WGS datasets by `MHCfinder` cannot guarantee the exact number of MHC-I genes within a particular species, it can make claims in studies of several species or taxanomic orders.

**Figure 6:**
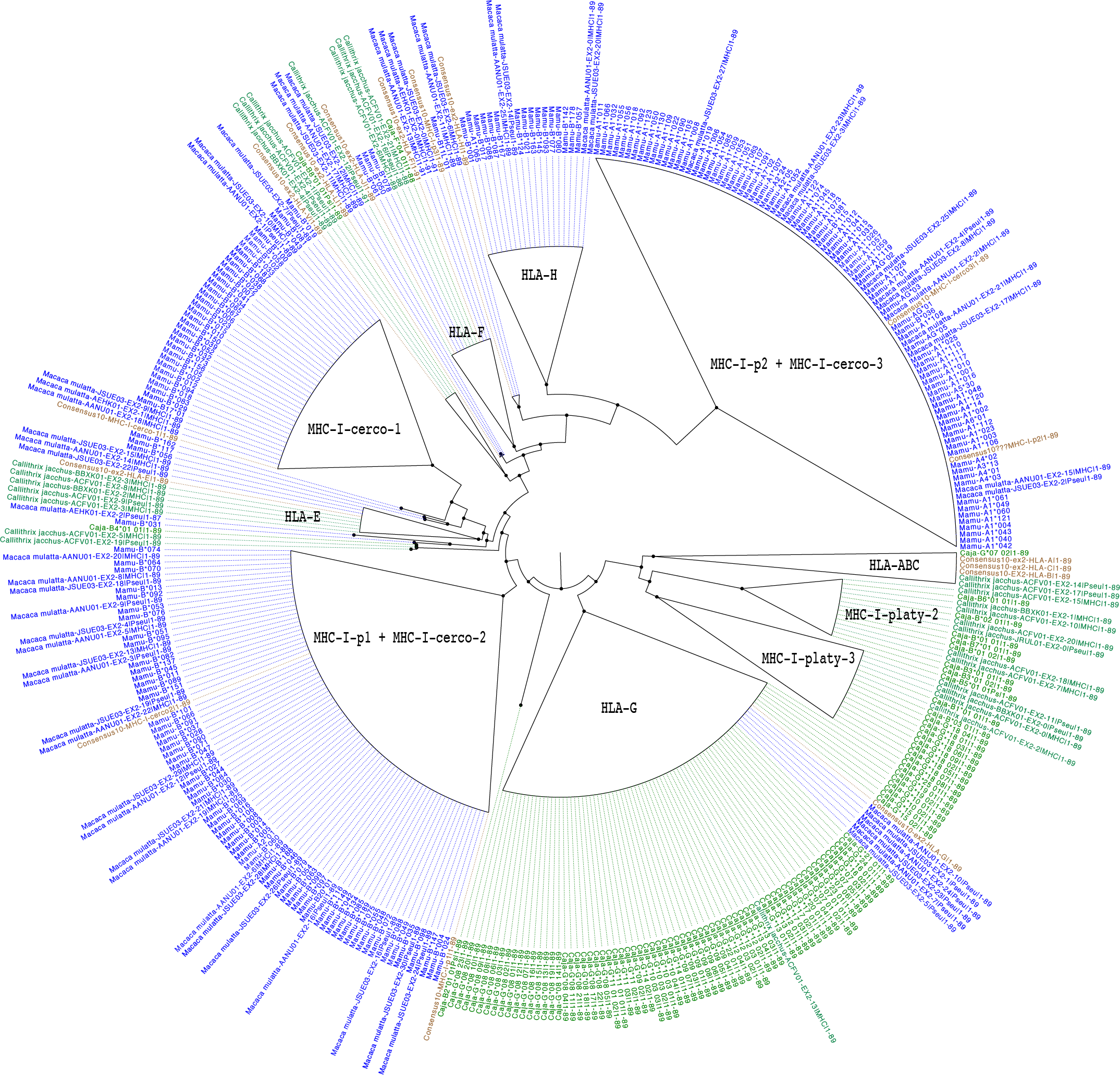
The phylogenetic tree using germline EX2 sequences from five WGS assemblies: two from *C. jacchus* and three from *M. mulatta*. Next, the allelic sequences of *M. mulatta* (blue) and *C jacchus* (green) obtained from the IDP-MHC database were aligned with the EX2 WGS sequences. Clades are collapsed to improve the visualization of the results. Also, consensus sequences (brown) were aligned with the EX2 sequences to identify clades.

For the first time, a large number of germline MHC-I gene sequences have been obtained across a wide range of primate species. Until now, sequences were obtained from the genomes or cDNA sequences of only few specific primates ((Heimbruch et al., 2015; Uda et al., 2005; Yan et al., 2013)). Moreover, this work can shed more light on previous studies of nonhuman primates that have identified expressed MHC-I alleles, attempting to classify these sequences without exact knowledge of the genes present in these species.

Given the ability to construct phylogenetic trees from the MHC-I exons of various species, it is possible to group the evolutionary origin of these genes and more precisely infer orthologs and paralogs. Here, sequences of exon EX2 were used to construct such trees; EX2 being the most discriminative constituent of the MHC-I gene, while EX3 and EX4 alone cannot resolve clades. The results indicate that the diversification of these genes has been driven by birth and death processes, thereby explaining the large number of pseudogenes.

The results presented here demonstrate that the classical HLA-A, -B and -C genes found in Hominids were generated recently, coinciding with the separation from Old World monkeys. In non-Hominid primates, orthologs to these genes can no longer be found, but instead there are paralogs that proceeded from a common ancestor. The sequences from Cercopithecus are practically absent, except for one sequence from *M. mulatta*, which is of interest since it corresponds to a gene having a large amount of allelic variability. Similar to these genes, which are quasi-specific to Hominids, there are other genes that are specific to Cercopithecus and Platyrrhini. Taken together, data from this study provides evidence of rapid birth and death processes. The absence of orthologs to HLA-A, -B and C in Cercopithecus, and to a lesser extent in Platyrrhini, may explain the generation and/or expansion of other clades consisting of genes that have the functions of the classical MHC-I. Confirmation of this hypothesis is confirmed in several cases, as seen in *M. Mulatta*, for which an allelic variation is seen that is not present in Hominids, while in *C. jacchus* a gene was identified belonging to the HLA-G that is responsible for most of the allelic variation.

The presence of orthologs to the nonclassical genes (HLA-E, HLA-F, HLA-G and HLA-H) in non Hominid primates indicates a previous origin prior to the separation from New World monkeys. In Prosimians, no orthologs were found, thereby raising the question whether the separation between the classical and nonclassical MHC-I is general to all mammals or specific to Platyrrhini and Catarrhini. Similar results are found by other authors (Piontkivska & Nei, 2003; Fukami-Kobayashi et al., 2005).

The birth and death processes observed in the data are similar to that which occurs in the variable (V) regions of immunoglobulin (IG) and T-cell receptor (TCR) genes. In the case of MHC-I, the situation is more complex because a viable replication process involves at least six separate exons. The birth and death process must be related to the adaptability and survival of the species. This evolutionary mechanism creates a greater genetic variability in genes that must present antigen, as well as in those that must recognize antigen. Nonetheless, it is still unknown why some species possess more V genes and/or MHC-I genes than others, or whether the absolute number of such genes provides the species with immunological advantages. More studies are needed to clarify whether correlaciones exists between the number of these genes and to establish if coevolution processes are at play.

## References

Birch, J., Murphy, L., MacHugh, N. D., & Ellis, S. A. (2006). Generation and maintenance of diversity in the cattle mhc class i region. Immunogenetics, 58, 670–679.

Breiman, L. (2001). Random forests. Mach. Learn., 45, 5–32. doi:10.1023/A:1010933404324.

Cao, Y., Fan, J., Li, A., Liu, H., Li, L., Zhang, C., Zeng, L., & Sun, Z. (2015). Identification of mhc i class genes in two platyrrhini species. Am J Primatol., 77, 527–34. doi:10.1002/ajp.22372.

Castro, M., Morales, P., Fern´andez-Soria, V., Suarez, B., Recio, M., Alvarez, M., Martín-Villa, M., & Arnaiz-Villena, A. (1996). Allelic diversity at the primate mhc-g locus: exon 3 bears stop codons in all cercopithecinae sequences. Immunogenetics, 43, 327–36.

Cock, P., Antao, T., Chang, J., Chapman, B., Cox, C., Dalke, A., Friedberg, I., Hamelryck, T., Kauff, F., Wilczynski, B., & de Hoon, M. (2009). Biopython: freely available python tools for computational molecular biology and bioinformatics. Bioinformatics, 25, 1422–3. doi:10.1093/bioinformatics/btp163.

Daza-Vamenta, R., Glusman, G., Rowen, L., Guthrie, B., & Geraghty, D. (2004). Genetic divergence of the rhesus macaque major histocompatibility complex. Genome Res., 14, 1501–15.

Djurisic, S., & Hviid, T. (2014). Hla class ib molecules and immune cells in pregnancy and preeclampsia. Front Immunol., 5, 652. doi:10.3389/fimmu.2014.00652.

Fukami-Kobayashi, K., Shiina, T., Anzai, T., Sano, K., Yamazaki, M., Inoko, H., & Tateno, Y. (2005). Genomic evolution of mhc class i region in primates. PNAS, 102, 9230–4. doi:10.1073/pnas.0500770102.

Garcia, K., Adams, J., Feng, D., & Ely, L. (2009). The molecular basis of tcr germline bias for mhc is surprisingly simple. Nat Immunol., 10, 143–7. doi:10.1038/ni.f.219.

Grimsley, C., Mather, K. A., & Ober, C. (1998). Hla-h: a pseudogene with increased variation due to balancing selection at neighboring loci. Molecular Biology and Evolution, 15, 1581–1588.

de Groot, N., Blokhuis, J., Otting, N., Doxiadis, G., & Bontrop, R. (2015). Co-evolution of the mhc class i and kir gene families in rhesus macaques: ancestry and plasticity. Immunol Rev., 267, 228–45. doi:10.1111/imr.12313.

de Groot, N., Otting, N., Robinson, J., Blancher, A., Lafont, B., Marsh, S., O’Connor, D., Shiina, T., Walter, L., Watkins, D., & Bontrop, R. (2012). Nomenclature report on the major histocompatibility complex genes and alleles of great ape, old and new world monkey species. Immunogenetics, 64, 615–31. doi:10.1007/s00251-012-0617-1.

Hedges, S., Dudley, J., & Kumar, S. (2006). Timetree: A public knowledge-base of divergence times among organisms. Bioinformatics, 22, 2971–2972.

Heimbruch, K. E., Karl, J. A., Wiseman, R. W., Dudley, D. M., Johnson, Z., Kaur, A., & O’Connor, D. H. (2015). Novel mhc class i full-length allele and haplotype characterization in sooty mangabeys. Immunogenetics, 67, 437–445.

Heinrichs, H., & Orr, H. (1990). Hla non-a,b,c class i genes: their structure and expression. Immunol Res., 9, 265–74.

Horton, R., Wilming, L., Rand, V., Lovering, R., Bruford, E., Khodiyar, V., Lush, M., Povey, S., Talbot, C. J., Wright, M., Wain, H., Trowsdale, J., Ziegler, A., & Beck, S. (2004). Gene map of the extended human mhc. Nat Rev Genet., 5, 889–99. doi:10.1038/nrg1489.

Kono, A., Brameier, M., Roos, C., Suzuki, S., Shigenari, A., Kametani, Y., Kitaura, K., Matsutani, T., Suzuki, R., Inoko, H., Walter, L., & Shiina, T. (2014). Genomic sequence analysis of the mhc class i g/f segment in common marmoset (callithrix jacchus). J Immunol., 192, 3239–46. doi:10.4049/jimmunol.1302745.

Lee, N Ishitani, A., & Geraghty, D. (2010). Hla-f is a surface marker on activated lymphocytes. Eur J Immunol., 40, 2308–18.

Lefranc, M., Duprat, E., Kaas, Q., Tranne, M., Thiriot, A., & Lefranc, G. (2005). Imgt unique numbering for mhc groove g-domain and mhc superfamily (mhcsf) g-like-domain. Dev Comp Immunol., 29, 917–38. doi:10.1016/j.dci.2005.03.003.

Liu, Y., Li, A., Wang, X., Sui, L., Li, M., Zhao, Y., Liu, B., Zeng, L., & Sun, Z. (2013). Mamu-b genes and their allelic repertoires in different populations of chinese-origin rhesus macaques. Immunogenetics, 65, 273–80. doi:10.1007/s00251-012-0673-6.

Lynge-Nilsson, L., Djurisic, S., & Hviid, T. (2014). Controlling the immunological crosstalk during conception and pregnancy: Hla-g in reproduction. Front Immunol., 5, 198. doi:10.3389/fimmu.2014.00198.

Moscoso, J., Serrano-Vela, J., Pacheco, R., & Arnaiz-Villena, A. (2006). Hla-g, -e and -f: allelism, function and evolution. Transpl Immunol., 17, 61–4. doi:10.1016/j.trim.2006.09.010.

Pedregosa, F., Varoquaux, G., Gramfort, A., Michel, V., Thirion, B., Grisel, O., Blondel, M., Prettenhofer, P., Weiss, R., Dubourg, V., Vanderplas, J., Passos, A., Cournapeau, D., Brucher, M., Perrot, M., & Duchesnay, E. (2011). Scikit-learn: Machine learning in Python. Journal of Machine Learning Research, 12, 2825–2830.

Perelman, P., Johnson, W., Roos, C., Seu´anez, H., Horvath, J., Moreira, M., Kessing, B., Pontius, J., Roelke, M., Rumpler, Y., Schneider, M., Silva, A., O’Brien, S., & Pecon-Slattery, J. (2011). A molecular phylogeny of living primates. PLoS Genet., 7, e1001342. doi:10.1371/journal.pgen.1001342.

Piontkivska, H., & Nei, M. (2003). Birth-and-death evolution in primate mhc class i genes: divergence time estimates. Molecular biology and evolution, 20, 601–609.

Price, M. N., Dehal, P. S., & Arkin, A. P. (2010). Fast-tree 2–approximately maximum-likelihood trees for large alignments. PloS one, 5, e9490.

Robinson, J., Halliwell, J., McWilliam, H., Lopez, R., & Marsh, S. (2013). Ipd -the immuno polymorphism database. Nucleic Acids Research, 41, D1234–40.

Robinson, J., Mistry, K., McWilliam, H., Lopez, R., Parham, P., & Marsh, S. (2011). The imgt/hla database. Nucleic Acids Res., 32, D1171–6. doi:10.1093/nar/gkq998.

Rogers, J., & Gibbs, R. (2014). Comparative primate genomics: emerging patterns of genome content and dynamics. Nat Rev Genet., 15, 347–59. doi:10.1038/nrg3707.

Uda, A., Tanabayashi, K., Fujita, O., Hotta, A., Terao, K., & Yamada, A. (2005). Identification of the mhc class ib locus in cynomolgus monkeys. Immunogenetics, 57, 189–197.

van der Wiel, M., Otting, N., de Groot, N., Doxiadis, G., & Bontrop, R. (2013). The repertoire of mhc class i genes in the common marmoset: evidence for functional plasticity. Immunogenetics, 65, 841–9. doi:10.1007/s00251-013-0732-7.

Wilming, L. G., Hart, E. A., Coggill, P. C., Horton, R., Gilbert, J. G., Clee, C., Jones, M., Lloyd, C., Palmer, S., Sims, S. et al. (2013). Sequencing and comparative analysis of the gorilla mhc genomic sequence. Database, 2013, bat011.

Yan, X., Li, A., Zeng, L., Cao, Y., He, J., Lv, L., Sui, L., Ye, H., Fan, J., Cui, X. et al. (2013). Identification of mhc class i sequences in four species of macaca of china. Immunogenetics, 65, 851–859.

